# Automated microuidic cell culture of stem cell derived dopaminergic neurons in Parkinson’s disease

**DOI:** 10.1101/209957

**Authors:** Khalid I.W. Kane, Edinson Lucumi Moreno, Siham Hachi, Moriz Walter, Javier Jarazo, Thomas Hankemeier, Paul Vulto, Jens Schwamborn, Martin Thoma, Ronan M.T. Fleming

**Affiliations:** Luxembourg Centre for Systems Biomedicine, University of Luxembourg, 7 avenue des Hauts-Fourneaux, L-4362 Esch-sur-Alzette, Grand-Duchy of Luxembourg.; Fraunhofer Institute for Manufacturing Engineering and Automation IPA, Stuttgart, Germany.; Analytical BioSciences Division, Leiden Academic Centre for Drug Research, Leiden University, Einsteinweg 55, 2333CC Leiden, The Netherlands; Mimetas B.V, PO Box 11002, 2301EA Leiden, The Netherlands.

## Abstract

Parkinson’s disease is a slowly progressive neurodegenerative disease characterised by dysfunction and death of selectively vulnerable midbrain dopaminergic neurons leading mainly to motor dysfunction, but also other non-motor symptoms. The development of human in vitro cellular models with similar phenotypic characteristics to selectively vulnerable neurons is a major challenge in Parkinson’s disease research. We constructed a fully automated cell culture platform optimised for long-term maintenance and monitoring of induced pluripotent stem cell derived neurons in three dimensional microfluidic cell culture devices. The system can be flexibly adapted to various experimental protocols and features time-lapse imaging microscopy for quality control and electrophysiology monitoring to assess neuronal activity. Using this system, we continuously monitored the differentiation of Parkinson’s disease patient derived human neuroepithelial stem cells into midbrain specific dopaminergic neurons. Calcium imaging confirmed the electrophysiological activity of differentiated neurons and immunostaining confirmed the efficiency of the differentiation protocol. This system is the first example of a fully automated Organ-on-a-Chip culture and enables a versatile array of in vitro experiments for patient-specific disease modelling.

## Introduction

Laboratory automation is becoming increasingly prevalent in the life sciences [1, 2]. Automated cell culture has the potential to increase the quantity and the quality of experiments that can be completed in parallel and enables long-term cell culture maintenance with reduced manual labour [3]. Once an automated protocol is established, a robot can operate continuously without fatigue and with the same consistency and accuracy [2]. Likewise, once established an automated imaging system can take repeated measurements over a long period without intervention [4]. The combination of robotic cell culture and automated imaging has a wide range of biological applications. A leading example is their use to distinguish causation from correlation in the pathogenesis of neurodegenerative diseases by longitudinal measurement of human *in vitro* disease models [5]. Laboratory automation requires precise specification of, and enables fine control over, many experimental protocol parameters, such as dispensing speed, cell culture conditions, fluid temperature and measurements. This enhances experimental reproducibility by reducing variance between replicates [6]. *In vitro* cell culture automation facilitates faithful replication of certain *in vivo* physiological conditions as it enables quantitative control over key experimental parameters, e.g., perfusion rate [7]. This increases the validity of employing an *in vitro* model to represent an *in vivo* system, in health or disease, thereby accelerating biomedical research.

During manual cell culture, procedures involving liquid handling, such as dispensing media, aspiring media, and movement of liquid samples between containers, are essential to all protocols. Therefore, when a cell culture protocol is automated, a liquid-handler and a robot for transposition receptacles are two of the most important devices. There are two types of technologies used in liquid-handler: *contact* and *non-contact dispensing* [2]. To dispense a precise volume, contact dispensing requires the head of the tip holding the fluid to touch the bottom of the substrate, for instance the bottom of a well, or to touch the liquid surface. Non-contact dispensing does not require any contact between the tip and the substrate, or liquid surface, for liquid release. Dispensing can require the handling of very small volumes, as low as a few nano-litres, so the technological advances in liquid handlers have focused more on dispensing than on aspiration [2]. Low volume dispensing and aspiration are especially required for microfluidic cell culture [7]. The robot to move receptacles can be a robotic arm or a gantry robot with a gripper for receptacles [2]. A gantry robot only moves in Cartesian coordinates where the three principal axes of control have linear actuators.

The choice of devices used in laboratory automation should be based on their intended uses, flexibility, purchase costs and maintenance costs. Selecting the components of an automated plant usually entails having to purchase devices from different manufacturers as no single firm supplies all of the devices that might be required to automate a laboratory protocol. Therefore, all of the components must be amenable to software integration in order to be able to function as a single autonomous plant. *Computer scripting* achieves integration by assigning a master software that communicates directly with all devices [8]. In this approach, assuming that all the devices are able to send and receive commands, a communication protocol must be implemented that is compatible with each individual device. However, this approach requires the master device software to recognise every other device using an idiosyncratic communication protocol. This approach can be a very expensive and challenging to implement. Alternatively, *Standardisation in Laboratory Automation* (SiLA, http://www.sila-standard.org/) is a consistent and efficiently extensible approach for integration of laboratory automation devices based on a standard protocol specification for exchanging structured information in a client-server model of communication. Furthermore, SiLA defines over 30 standard device classes used in the field of life sciences, including incubators, microscopes, de-lidders and liquid handlers [9]. For each device class, a list of required and optional functions are proposed to standardise the software communication within a laboratory automation plant. This approach standardises the communication between all of the devices of a plant, regardless of the manufacturer, and a SiLA compatible process management software can then be used to control each SiLA compatible device without any modification.

Parkinson’s disease is the second most common neurodegenerative disease with more than 10 million people affected worldwide [10, 11]. Parkinson’s disease is characterised by cell death in selectively vulnerable parts of the nervous system [10]. These neuronal losses include cholinergic neurons in the pedunculopontine nucleus, noradrenergic neurons in the locus coeruleus and dopaminergic neurons from the substantia nigra pars compacta [12, 13]. Dopaminergic neurons play a critical role in brain function by releasing a neurotransmitter called dopamine [14–16]. The loss of dopaminergic neurons is the main reason behind the motor symptoms (rigidity, tremors and postural instability) of Parkinson’s disease patients [17]. The study of Parkinson’s disease at the cellular level has been facilitated by the use of *induced pluripotent stem cells* (iPSCs) technology [18]. iPSCs are embryonic-like stem cells that have been derived from somatic cells, skin fibroblast, via reprogramming [19]. Reinhardt et al. [20] developed a protocol to generate human neuroepithelial stem cells (hNESCs) from iPSCs. These hNESCs can in turn be differentiated into many neuronal cell types, including midbrain-specific dopaminergic neurons, critical to the *in vitro* modelling of Parkinson’s disease pathogenesis.

Microfluidic cell culture concerns the design and implementation of devices and protocols for the culture, maintenance and perturbation of cells in micro-scale fluid volumes. The reasons behind the popularity of microfluidic cell culture are both economic and scientific. Cell culture reagents are expensive, and the amounts used in microfluidic cell cultures are much less than in macroscopic cell culture [21, 22]. Microfluidic cell culture also has the potential to lower the ratio of extracellular to intracellular fluid volumes, thereby decreasing the temporal lag in extracellular response to molecules transported across cell membranes, e.g., in exometabolomic analyses [23–25]. With the advent of Organ-on-a-Chip technology [26] microfluidic cell culture has developed tremendously and includes examples of perfusion culture, co -culture and three dimensional cell cultures [27–29]. Moreover, miniaturisation enables multiple experimental replicates within a geometrically confined experimental footprint. Thus far, no examples are known of Organ-on-a-Chip operation in an automated setting although few hold the promise to do so [30]. Even though the combination of automation, microfluidics and cell culture technologies allows the screening of multiple environmental conditions in parallel [31, 32] as well as enabling regular live cell culture monitoring [33, 34], at a temporal resolution impractically in a manual setting. Therefore, laboratory automation technology is key to unleash the full potential of microfluidic cell culture. We previously developed a microfluidic titer plate for three dimensional microfluidic cell culture, called OrganoPlate [28]. Subsequently, we implemented the differentiation of hNESCs into three dimensional networks of electrophysiologically active dopaminergic neurons into the OrganoPlate [35]. The microfluidic titer plate was designed for compatibility with laboratory automation, but this has yet to be exploited. The manual culture of human pluripotent stem cell derived cells within the microfluidic titer plate has also been established, but the potential for automation has also not yet been exploited.

Herein, we report the integration of developmental biology, microfluidic cell culture and laboratory automation technology to generate a flexible, fully automated, enclosed microfluidic and macroscopic cell culture observatory, termed the *Pelican*. We elaborate on each device in the Pelican, as well as the SiLA software integration approach used to realise a fully automated system. We illustrate the functionality of the Pelican for automated cell culture and differentiation of human neuroepithelial stem cells into dopaminergic neurons within a three-dimensional microfluidic device [28]. We monitored the health of the cells throughout the experiment with an automated image acquisition pipeline. After 24 days in culture, we assessed the outcome by characterising known features of dopaminergic neurons by calcium imaging and immunofluorescence assays. Three dimensional imaging revealed mature and interconnected neuronal populations within microfluidic cell culture chips. The Pelican is a modular automation system, compatible with implementation of a variety of automation platforms, where cost-effective flexibility is maximised to allow for replacement or further expansion of platforms by integration of new devices. Microfluidic cell culture has already been integrated with IPSC technology [35]. Our work integrates a fully automated Organ-on-a-Chip culture.

## Results

### System Construction and Design

Figure 1A illustrates the fully automated cell culture enclosure (see Supplementary Figs S1 and S2 online), constructed and assembled according to the hardware design in Figure 6 and the plant control architecture in Figure 1B. The different devices and their use in the automated system are detailed in the Supplementary Experimental Procedures. In addition, a user manual that is continuously maintained, and that details the procedure for operation of the Pelican Cell culture Observatory is available upon request. This document detailed not only how to operate the system, but also how to extend the current setup to include more devices, and how to integrate the said devices in terms of hardware and software development.

**Figure 1:**
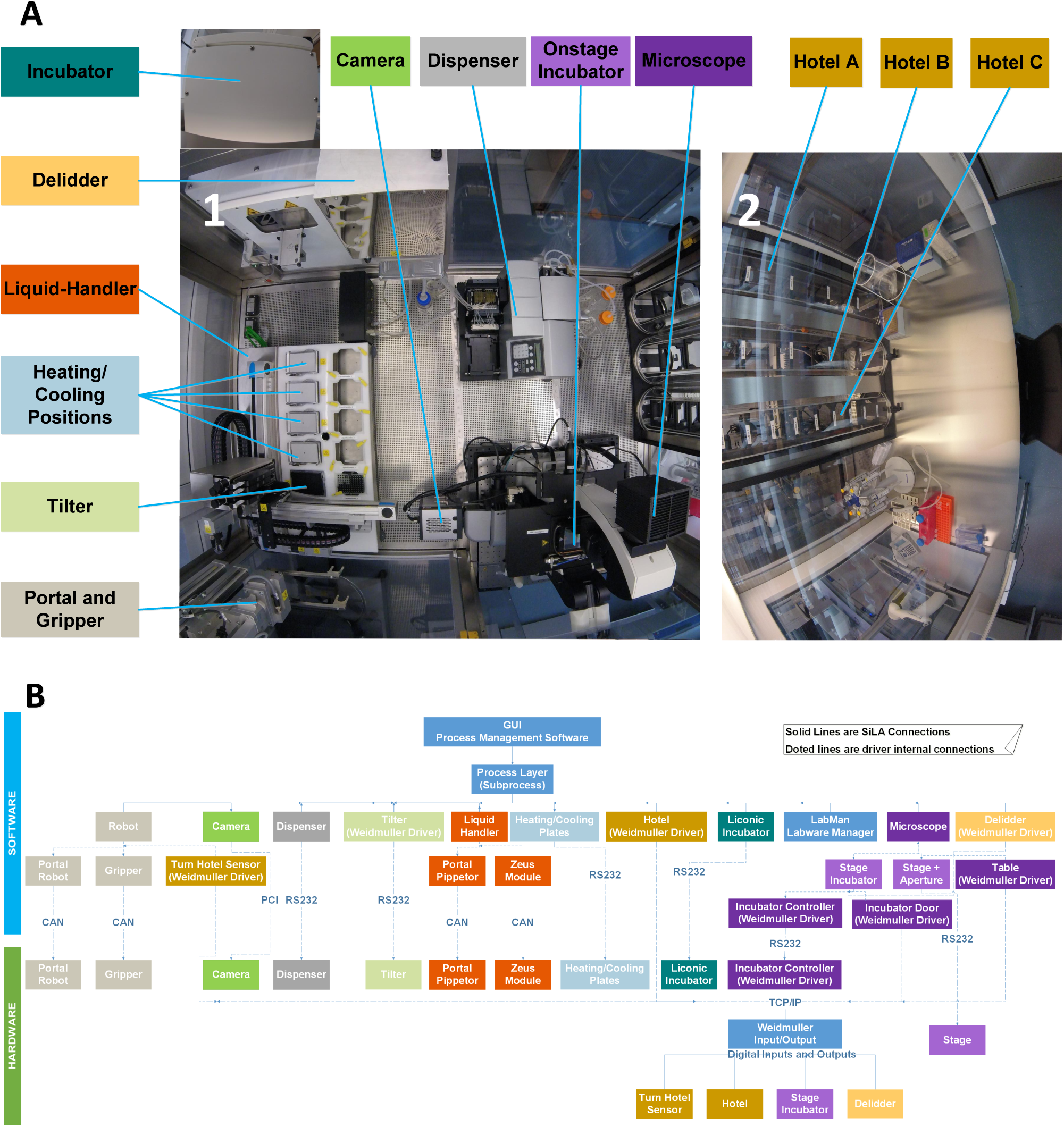
Pelican automated cell culture observatory. (A) Top view inside the Pelican automation worksta-tion without housing: (1) Wide angle lens image of the automated enclosure. (2) Wide angle lens image of the adjacent manual cell culture bench. (B) Pelican hardware and software control architecture. Standard on arrow = type of physical connection between a device and the computer. CAN = Controller Area Network. PCI = Peri-pheral Component Interconnect. RS232 = Recommend Standard number 232. TCP/IP = Transmission Control Protocol/Internet Protocol. The colour codes of the devices labels in (A) and (B) match.

### Automating the differentiation of human neuroepithelial stem cell into dopaminergic neurons

The day after loading the gel-embedded hNESCs into the culture lane of the OrganoPlate, an *automated image acquisition protocol* was executed through LACS to qualitatively assess the health of the cell culture. The automated image acquisition protocol (Figure 3) consisted of first setting the environmental condition of the onstage incubator to 5% CO_2_ and to a temperature of 37°*C*. Second, the robotic arm moved the plate from the storage incubator to the microscope. Third, the microscope scanned through all the observation windows of the OrganoPlate, and the camera took an image of each observation window. Fourth, the plate was transported back to the incubator by the robotic arm. Immediately after the automated image acquisition, the *automated differentiation of hNESC into dopaminergic neurons protocol* (Figure 2) was executed through LACS. This protocol consisted of first instructing the robotic arm to move the plate from the storage incubator to the de-lidder to remove the lid from the plate, before moving it to the liquid dispenser. Second, the dispenser aspirated media from each medium inlet and outlet well. Third, the dispenser replenished the media in each medium inlet and outlet well respectively. Fourth, the robotic arm moved the plate to the de-lidder to put the lid back before placing the plate inside the storage incubator (Supplementary Video 1). These two protocols were run in this order every two days for 24 days.

**Figure 2:**
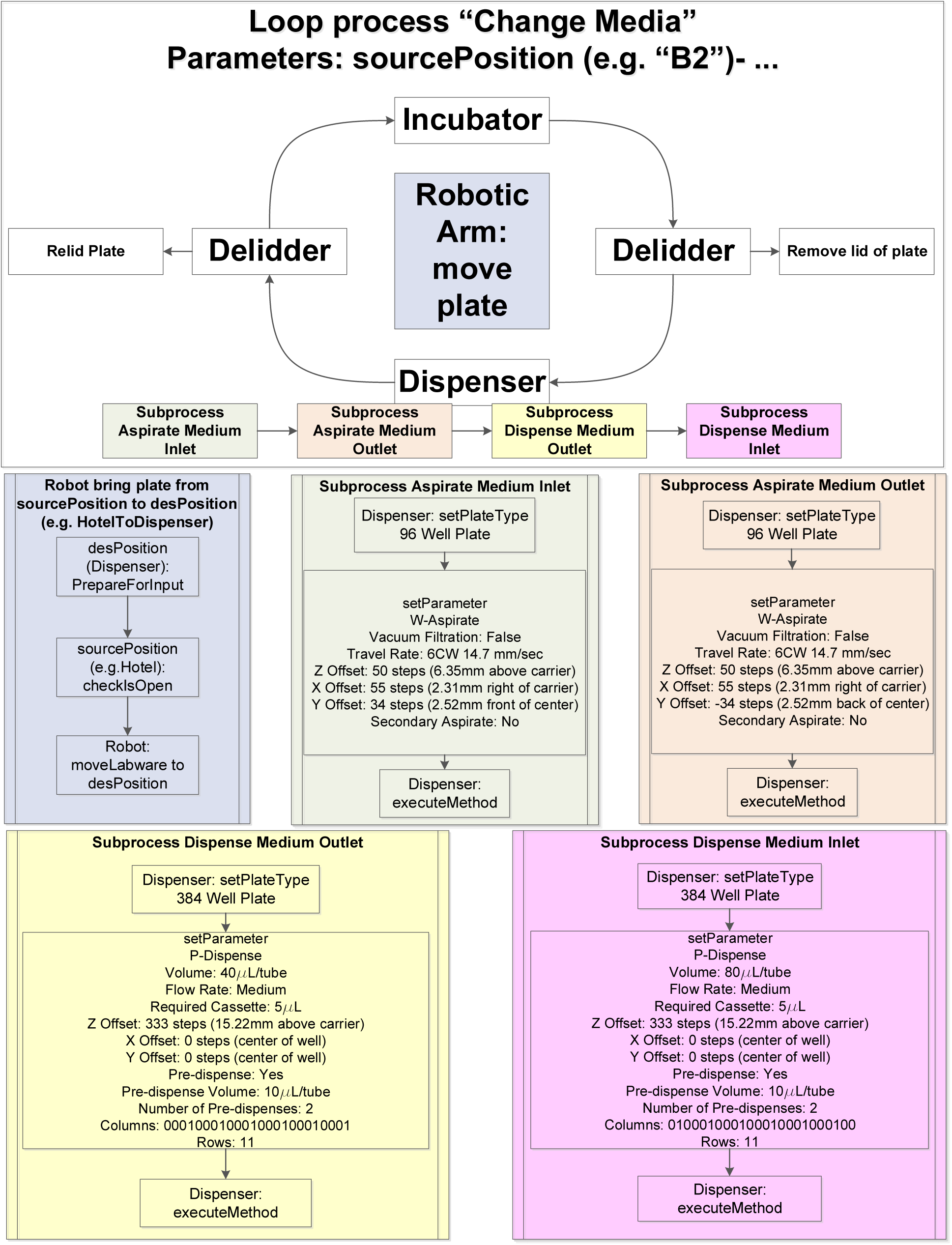
Automated media change pipeline. W = Washer, P = Peristaltic, 0 = Method not applied to column, 1 = Method applied to column, −1 = search position. Curved connection = Plate movement. Straight connection = Device specific method.

**Figure 3:**
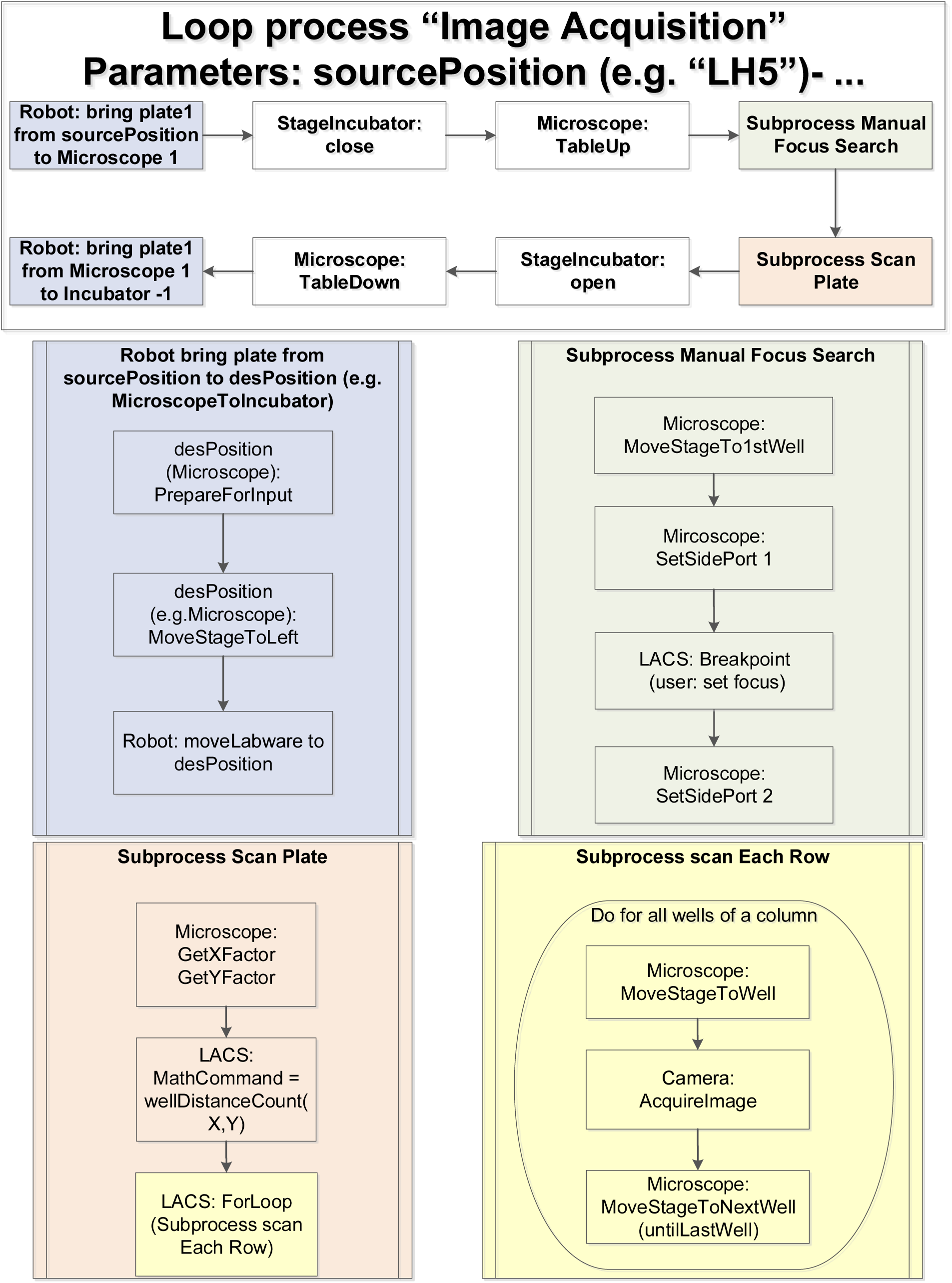
Automated image acquisition pipeline. Abbreviations as in Figure 2.

### Human neuroepithelial stem cell culture differentiation

We utilised the Pelican and the existing protocol described by Reinhardt et al and Lucumi et al [20, 35] to fully automate the differentiation of hNESC into dopaminergic neurons inside a stratified three-dimensional microfluidic device. After manually seeding hNESC in an OrganoPlate, a three dimensional microfluidic cell culture device compatible with laboratory automation the hNESC were distributed in three dimensions within the culture lane, but also adjacent to the meniscus (Figure 4A). Thereafter the Pelican executed an automated differentiation protocol to start and maintain the differentiation process. Initially this required a liquid dispenser to aspirate the maintenance medium and dispense the differentiation medium with PMA, via a dispensing cassette, into inlet and outlet wells of the OrganoPlate. Thereafter it required incubation and regular media replenishment.

**Figure 4:**
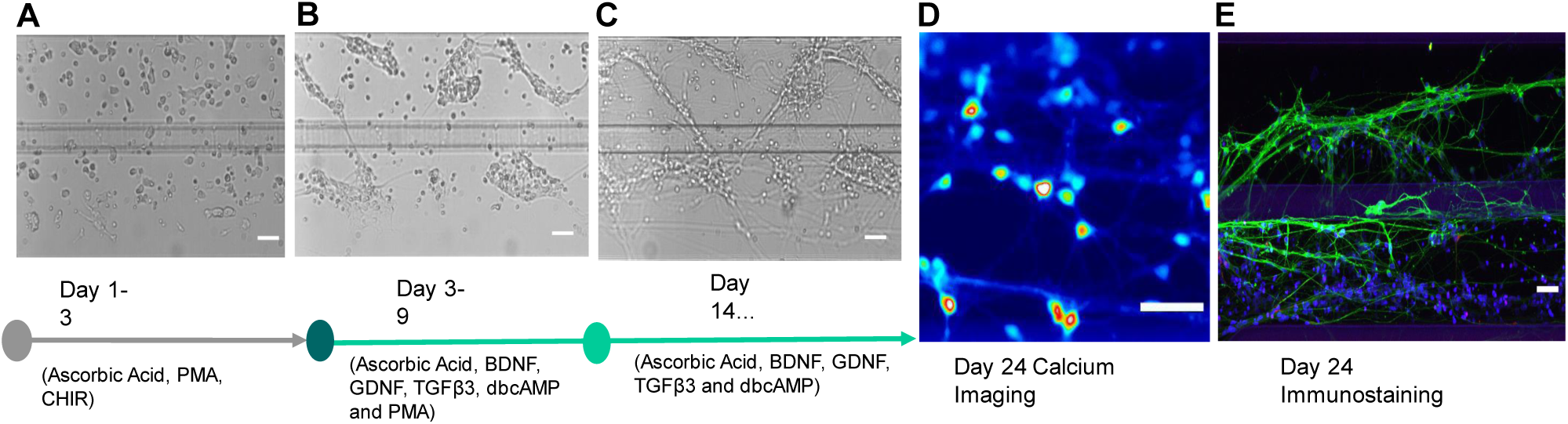
Dopaminergic neuronal differentiation. Bright field images of hNESCs with media components at (A) 1 day (B) 4 days and (C) 21 days after seeding. (D) Calcium imaging frame of a firing event taken at day 24 after seeding. (E) Immunofluorescence image illustrating the neuronal composition inside a culture chamber. All scale bars are 50 µm. Legend as in Figure 5.

During the differentiation, cells started to form aggregates, and morphological changes started to appear, such as acquisition of cellular polarity and projection of processes representative of neuronal morphology (Figure 4B). An automated protocol replaced the medium with PMA with medium without PMA, and differentiated neurons started a maturation process accompanied by acquisition of a more evident neuronal morphology (Figure 4C). Differentiated neurons projecting their processes can be observed in the culture lane and in the area occupied by the meniscus, located in part of the medium lane. Neuronal activity of differentiated cells was tested using calcium imaging (Figure 4D) and morphological and phenotypic characteristics like neuronal processes positive for TUBβIII (green) and the amount of tyrosine hydroxylase positive neurons (red) were characterised using an immunostaining assay (Figure 4E). A top view of the culture chamber of a chip with hNESC (Figure 5A), shows that cells were distributed in the entire culture chamber, Hoechst nuclear staining (blue). Furthermore, their processes have been projected in all directions of the culture lane, as well as on the meniscus part in the medium lane. It can also be seen that the neuronal networks and the level of connectivity of the differentiated neurons is homogeneous in the entire culture chamber, denoting an even effect of the differentiation protocol, as well as the efficiency of perfusion in the medium lane (Figure 5A).

**Figure 5:**
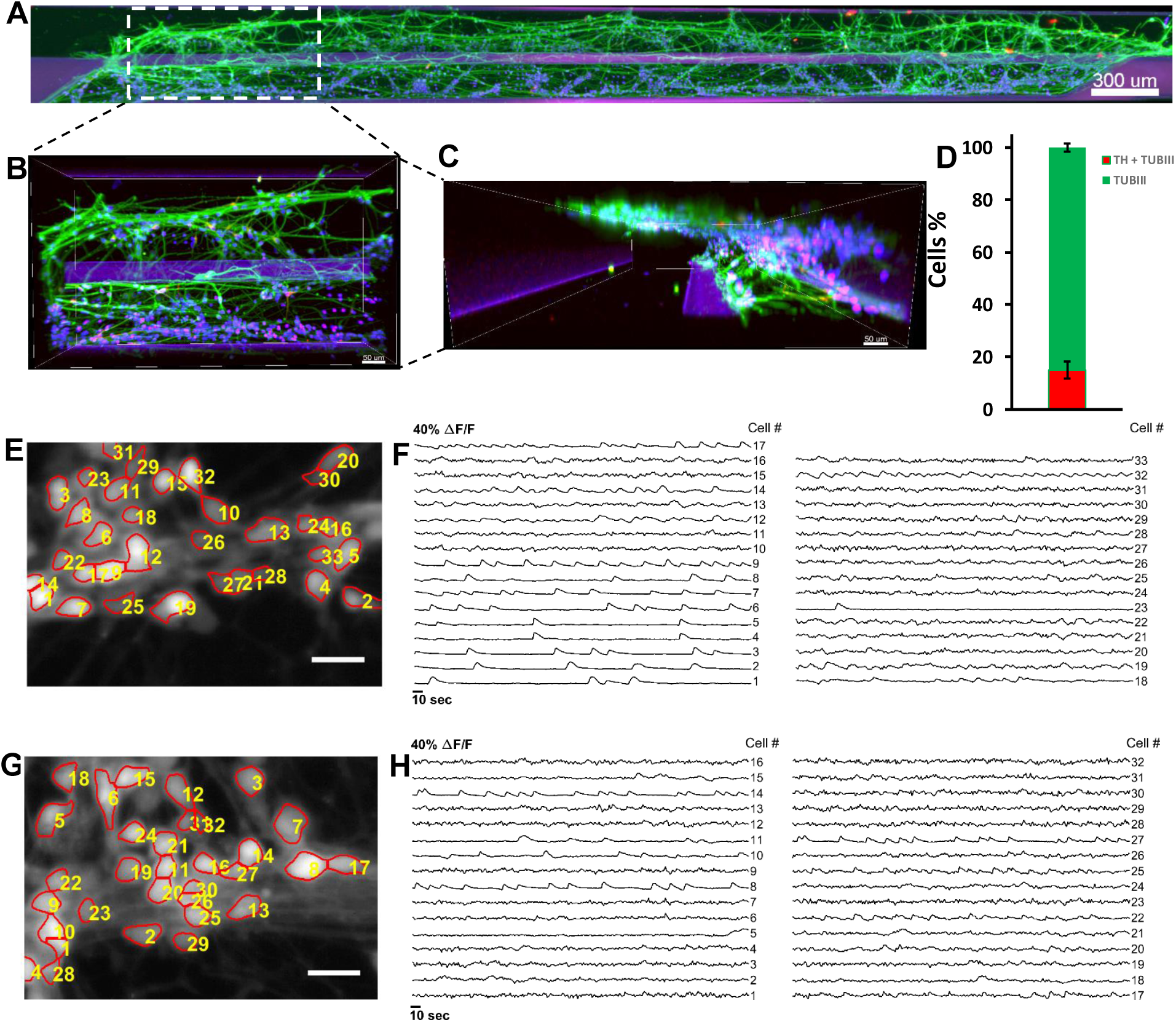
Immunostaining and automated cell segmentation of calcium time-series hNESC differentiated in chips of the OrganoPlate inside the Pelican. A) Top view of an entire chip in the OrganoPlate showing differentiated wild type neurons (K7 cell line) immunostained for nuclei with Hoechst (blue), TUBβIII (green) and tyrosine hydroxylase (red); scale bar 300µm. (B) Enlarge top (B) and front (C) views of selected area; scale bar 50µm. (D) Differentiation efficiency of neurons positive for TUBβIII and tyrosine hydroxylase. (E) Mean fluorescence frame of a calcium time-series of WT population with segmented regions of interest corresponding to individual neurons and their corresponding fluorescence traces (F). (G) Mean fluorescence frame of a calcium time-series of PINK1 mutants with segmented regions of interest corresponding to individual neurons and their corresponding fluorescence traces (H).

Figure 5B enlarges the view of a specific area of one chip with differentiated neurons from hNESC. Neurons positive for TUBβIII (green) and positive for tyrosine hydroxylase (red) are located in both culture and medium lanes. However, all cells appear to migrate towards the medium lane. This is confirmed in the front view of the enlarged area (Figure 5C), where the extent of the area occupied by the meniscus in the medium lane is clear, in addition to the degree of cells located on the meniscus on the medium lane. On average, the efficiency of differentiation for tyrosine hydroxylase positive neurons in 3 chips of the OrganoPlate with hNESC differentiated neurons was 15% of all neurons (Figure 5D), which is in accordance with values reported previously in analogous manual, microfluidic and macroscopic cell culture systems [20, 35].

After 24 days, we were able to obtain midbrain-like, mature dopaminergic neurons in 96 three-dimensional self-contained microfluidic chips. Regular pictures were taken during differentiation for quality control and morphological study (Figure 4A,B,C), and end point assays are represented in Figure 4D,E to illustrate the fate of the differentiation.

### Calcium imaging and immunostaining assays

To probe the neuronal activity of cells cultured in an OrganoPlate within the Pelican, we used Fluo-4-based calcium imaging. We acquired time-series of representative culture chambers of WT and PINK1 p.I368N-mutated populations at day 24 of differentiation (Supplementary Video 2). Analysis of those calcium imaging data revealed spontaneous neuronal activity in differentiated neurons control and PINK1 -mutant neurons in an OrganoPlate, within the Pelican. We detected individual cells by applying an automated cell segmentation algorithm [36] to the raw calcium imaging data. Figures 6E and 6G illustrate segmented mean fluorescence frames of representative culture chambers of an OrganoPlate with WT and PINK1-mutated neurons respectively. Fluorescence traces were then measured for each segmented cell to assess their activity (Figure 6F and H). Some of the fluorescence traces reveal calcium transients indicating neuronal firing events (Figure 6F e.g. signal #2, 3, 9 and Figure 6H e.g. signal #8, 14, 27). Fluorescence traces revealed different firing patterns of the differentiated neurons. Some of these traces exhibited regular firing patterns, as opposed to other ones, corresponding probably to dopaminergic neurons, similar to what has been previously reported by Lucumi et al. [35] in adjacent manual culture.

**Figure 6:**
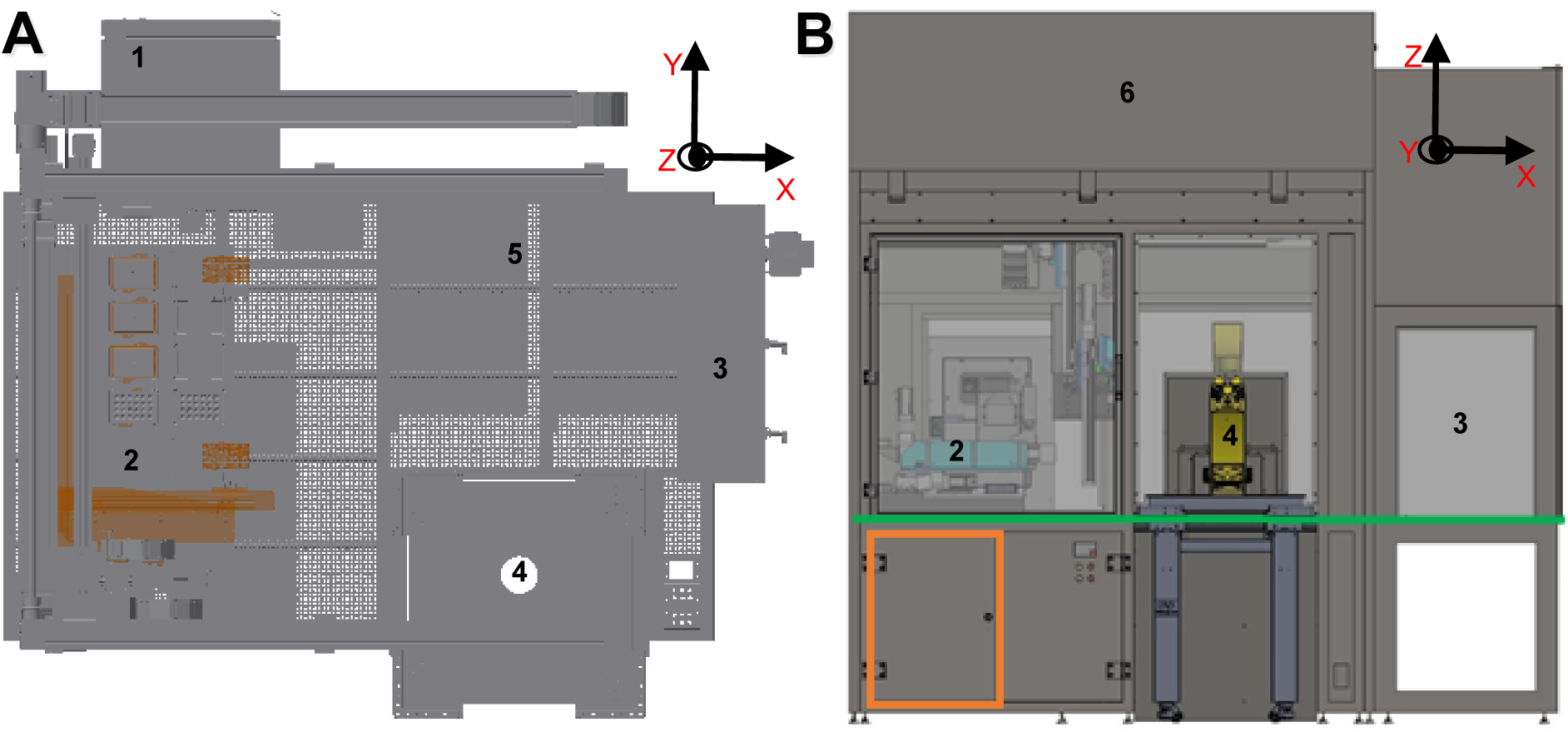
CAD drawing of automated workstation. (A) Top view inside the Pelican without housing. (B) Front view of the Pelican with housing. Yellow (imaging station), light blue (liquid handling station), green (level of stainless steel work surface) and orange (waste containers). Perforated surface=work surface. (1) Storage incubator. (2) Liquid-handler. (3) Hotels (manual working bench). (4) Position of microscopy station. (5) Position of liquid dispenser. (6) Automated work bench. Coordinates shown here are consistent with the remainder of the manuscript. See also Figures S1 and S2.

## Discussion

We assembled a fully automated cell culture observatory, termed the Pelican. It was optimised for the long-term cell culture maintenance of neurons inside three-dimensional microfluidic devices. The implementation followed a modular automation design, with generous space for further devices, and is flexible in terms of the automation platforms that can be implemented. For example, the system allows the automation of seeding, feeding and other cell culture processes that could initiate and maintain cell lines in any standard microtiter plate format. In this regard, we carefully selected the robotic system, the pipetting technologies and the techniques to integrate all the components of the system into a single plant.

In the Pelican, a four-axis (X-Y-Z-θ) gantry robot in combination with a gripper, that allows many types of substrates to be handled and services all devices with substrate from above. The robot was attached to the top of the stainless steel frame support inside the housing through rails that allow the movement of the robotic arm along three axes. This maximised the modular capacity of the Pelican because the rails determine the robotically *useful space* of the system, in contrast to other automated systems with a fixed rotating robot arm, which often has limited reach. The useful space is the space that is available to potentially hold new devices, adding to the functions of the automated platform.

In the Pelican, a ZEUS pipetting module (Hamilton Inc.) with disposable tips combines the precision of contact dispensing with the versatility of non-contact dispensing. In addition, a liquid dispenser with only non-contact dispensing was also implemented in the system. The liquid dispenser is less precise than the liquid-handler. However, it is much faster as it can handle up to 96 wells per step compared to 4 wells per step for the pipetting modules. Despite its shortcomings, the contact dispensing function of the ZEUS is especially useful for microfluidic cell culture where very small volumes must be dispensed. In addition, the contact dispensing helps to make sure that the dispensed media is bubble free. This is very important as with low flow rate non-contact dispensing, bubbles that arise can imped the flow of fresh media, which could ultimately starve the cells in a chip. Contact dispensing is very precise for dispensing small volumes. However, this precision is dependent on dispensing at an exact location, which is not always possible as the dispensing tip cannot always physically access the well to make contact with the liquid [1].

With contact dispensing, well cross-contamination is a risk as there is direct contact between the tip and the liquid in the destination well. Therefore, a cleaning protocol was implemented after each dispensing step. This does not promote speed and high-throughput capabilities so sought after in laboratory automation, however, the contact dispensing was only utilised for the initial loading of media to avoid the introduction of bubbles in the dry medium lane. Non-contact dispensing does not require any contact between the dispensing tip and the liquid. This helps to avoid cross-contamination, and promotes the integrity of the well. Non-contact dispensing is very popular in laboratory automation because it is versatile, and it is easy to dispense to any area of a well regardless of geometry such as undercuts, so long as there is an opening on the well [37]. As a result, the non-contact dispensing methodology was utilised for all subsequent media changes.

Many methods have been developed over the years to address the issue of standardisation and easy integration of new devices into existing laboratory automated plants [8, 9, 38–41]. In the Pelican, Standardisation in Laboratory Automation (SiLA) was chosen for the integration of the devices. SiLA standardises the interfacing, integration and data representation in a simple single XML schema for all devices [9, 42]. SiLA standardises the communication between process management software and one or more devices. SiLA defines common commands per *device class* and the common *device states*. SiLA defines the device classes to achieve these functionalities such as an incubator and a liquid-handler. Each device class will have the required commands for the core functionalities as well as *optional commands* for extended functionalities that are not necessarily present in every device of the same class. In this regard, a SiLA standard common command dictionary was developed for every single device class where the commands such as setParameter and getParameter and the expected return are known. This allows a process management software to automatically generate the required commands for every device class. Each component of the Pelican was chosen based on cell culture needs without taking into account the manufacturer of each device. The integration of any new device would require sufficient space within the useable area and possibly the development of a SiLA driver (tutorial in user manual available upon request). The modular hardware and software integration flexibility is a key advantage of the Pelican design compared to other automated cell culture systems [43–48].

In order to demonstrate the biological utility of the Pelican, control and PINK1-mutant human neuroepithelial stem cell lines were automatically differentiated into midbrain specific dopaminergic neurons. Calcium imaging of spontaneously firing neurons, as well as immunostaining for neuronal markers demonstrate that neuroepithelial stem cells could be successfully maintained and were spontaneously active within the OrganoPlate inside the Pelican. Further analysis of fluorescence traces for additional cell lines with different genetic backgrounds would be necessary to quantify any difference in phenotypic characteristics of neuronal activity between control and PINK1 mutants neurons. The differentiated neurons can be maintained inside the OrganoPlate for at least 100 days. The Pelican demonstrates proof-of-concept for automated generation of personalised *in vitro* neuronal models from human neuroepithelial stem cells via a microfluidic cell culture approach.

The Pelican is designed for longitudinal analysis of many personalised cellular models exposed to a few perturbations, rather than single, end-point analysis of one cellular model exposed to a large number of perturbations, as for instance is the focus in high throughput drug screening. Therefore, we envisage that such automated system be applied to stratification of patients with complex diseases. For example, idiopathic Parkinson’s disease patient groups display statistically significant reduction in mitochondrial activity [49]. However, except for genetic analyses, due to the broad normal range of many clinically accessible and mitochondrial associated parameters, it is not currently possible to sub-classify individual Parkinson’s disease patient into those with and without overt mitochondrial dysfunction. Phenotypic deficiencies have been measured in patient-specific in vitro neuronal models of Parkinson’s disease that are otherwise inaccessible in a clinical environment.

We envisage the use of automated cell culture to enable comprehensive phenotyping of large, parallel sets of personalised, in vitro, midbrain-specific, dopaminergic neuronal models of Parkinson’s disease. By integrating the data generated with a generic mechanistic computational model of the underlying biochemical network of a dopaminergic neuron, each personalised computational model then becomes a coherent representation of our information about the cell autonomous characteristics of Parkinson’s disease in each patient. Such personalised computational models, and thereby the corresponding patients, are then amenable to stratification with a range of powerful stratification tools. Stratification based on personalised computational models of the personalised data is statistically superior to stratification based on the personalised data alone as the former explicitly incorporates the wealth of prior biochemical information known about midbrain dopaminergic neurons. The aetiopathogenic stratification of Parkinson’s disease patients, for instance into mitochondrial vs non-mitochondrial Parkinson’s disease is an important first step toward targeting the development of therapies toward the underlying dysfunction that is present. Ultimately this will accelerate the translation of basic biomedical knowledge from the laboratory to the therapies with clinical impact.

Increasing demands for reproducibility, parallelisation and longitudinal observations are driving cell culture research toward automation. We developed a novel automated cell culture observatory that enables long-term maintenance and longitudinal optical measurement of cellular parameters in Organ-on-a-Chip platforms. We demonstrate the use of this platform to successfully automate the generation of personalised in vitro neuronal models from human neuroepithelial stem cells. We demonstrate the feasibility of automated image acquisition on this platform and compatibility with different real-time and end-point assays. It is the first time that an Organ-on-a-Chip platform is applied in a fully automated setting. It holds great promise for patient stratification by enabling comprehensive phenotyping of large, parallel sets of personalised, in vitro, models of complex diseases.

## Methods

### System Construction and Design

The Pelican is composed of a sterile automation enclosure that abuts a sterile manual enclosure on one side and an incubator on another. The automation enclosure contains a set of devices that may physically communicate via a four-axis gantry robot within a customised housing support. The manual enclosure is a cell culture hood, adapted for restricted communication of material with the automation enclosure. The automation enclosure currently includes a de-lidder, eight-fold and 96fold parallel dispenser, three-axis fourfold liquid handling robot (pipettor) with disposable tips, confocal microscope, and camera. The assembly (Figure 6B), all of the components are described in the Supplementary Experimental Procedures.

#### System integration

Each of the devices of the Pelican are physically connected via Ethernet to a computer (Precision T7600, Dell Sa, Mamer, Luxembourg), according to the plant architecture illustrated in Figure 1B. The SiLA standard was consistently implemented for networking and integrating the devices and components. A driver development kit (DDK, Fraunhofer IPA, http://www.sila-standard.org/driver-information-platform/fraunhofer-ipa/sila-driver-development-kit/) enabled the development of SiLA compliant drivers for each of the devices, if it was not available from the manufacturer. Incorporating a driver and converting the commands occurred in *Laboratory Automation Control Suite (LACS)*. More information about LACS, and how to operate it is available on request.

In brief, LACS is a programming environment developed by Fraunhofer IPA for laboratory automation. LACS incorporates three software packages: LacsDriverCore, LACS graphical user interface (GUI) and LACS Config Editor. LacsDriverCore reads the SiLA drivers of the devices generated by the DDK, hence, it connects the devices to a computer. The LACS GUI is the process management software, the interface between a device and an operator (via the computer) through a loaded SiLA driver. LACS Config Editor is used to draft automated protocols. LACS does not make any process-specific decisions, nor does it analyse any data. In the event of an unexpected event (a device failure or any other error), the operator is always prompted to assess and rectify the error or the event.

The safety features around the housing, the hotels and the incubator of the Pelican (sensors to read the states of the components), and the operations of the de-lidder and the stage incubator were all controlled through a *digital and analogue logic* module (UR20-FBC-MOD 1334930000, Weidmuller GmbH & Co. KG, Germany). A digital and analogue logic device has a binary set; a binary input and binary output. It is usually used for a maintenance device or to control a device with a simple binary command such as for a valve ON/OFF and a light switch. A single weidmuller SiLA driver was installed to control all devices connected to one logic module.

A software called *labware manager* handles all positions inside the pelican plant and all substrates. In the first case it holds the information, which position is occupied and with which substrate and which free position could potentially hold which type of substrate. The labware manager knows each position and status as well as each substrate in the plant including position and type. All data are stored in an open source PostgreSQL database (*PostgreSQL*, https://www.postgresql.org/). Like the hardware devices, this virtual device has a SiLA communication interface and a Windows 7 graphical user interface to view the stored data.

### Microfluidic device

A 2-lane *OrganoPlate* (#*9603-200B*, Mimetas BV, Leiden, The Netherlands) consists of a stratified array of 96 microfluidic chips embedded in a customised 384-well microtiter plate format [28] (Figure 7). Each chip consists of a single microfluidic chip contained between two pieces of glass: a top plate with holes corresponding to the underside of selected wells, and a bottom plate. Each chip is connected through 4 neighbouring wells and 2 lanes: one *gel inlet well* for loading of gel-embedded cells into the culture lane, one *medium inlet well* connected to one *medium outlet well* though a medium lane where the flow of media is driven by a pressure drop between these aforementioned 2 wells, and one well used as an *observation window* for monitoring the quality of cells through an inverted microscope. The culture and medium lanes are separated by a *phaseguide*, preventing the gel-embedded cells from flowing into the medium lane forcing it to stay on the culture lane. A phaseguide is a patterned pinning barrier that controls the liquid-air interface by forcing it to align with the ridge, hence guiding the fluid flow into the needed lane [27].

**Figure 7:**
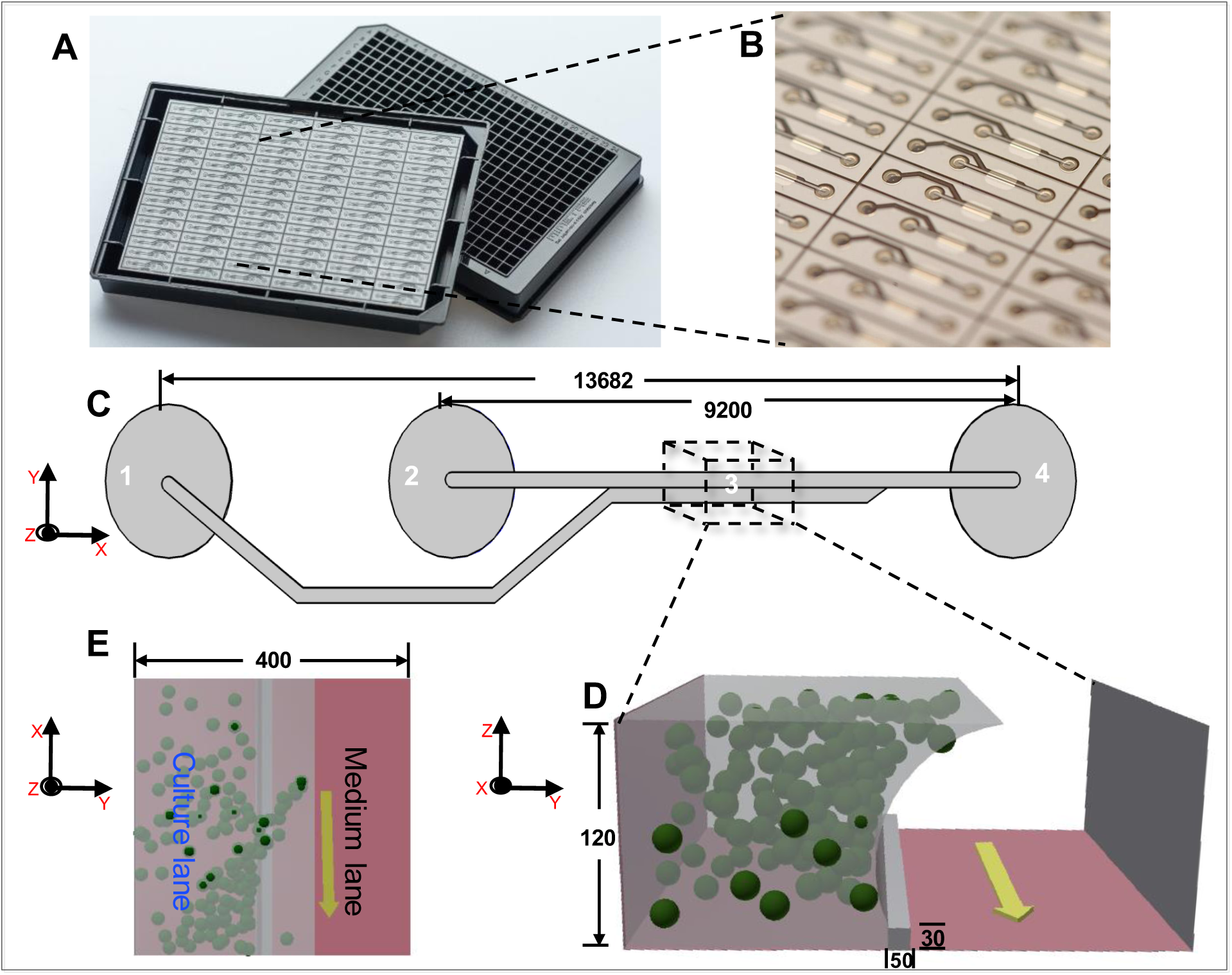
Microfluidic cell culture device: OrganoPlate. (A) Photograph of underside (left) and upper views (right) of an OrganoPlate. (B) bottom plate of OrganoPlate with selective chips. (C) A schematic of a single 2-lane chip; 1 = Gel inlet, 2 = Medium inlet, 3 = Observation window, 4 = Medium outlet. (D) A transverse section of a culture chamber showing the direction of the flow in the medium lane (yellow arrow), the phaseguide and the culture lane with suspended spheres representing cells embedded in Matrigel. (E) Top view of transverse section of a culture chamber with phaseguide, medium and culture lanes. All dimensions in µm.

### Cell culture

#### Human neuroepithelial stem cell culture

We used a human neuroepithelial stem cell line from a healthy donor (#hNESC K7) and a human neuroepithelial stem cell line derived from a patient carrying the Parkinson’s disease related mutation p.I368N in PINK1 (#40066C5N). These cells were maintained and differentiated into midbrain-specific dopaminergic neurons within an OrganoPlate, by automating an existing macroscopic cell culture protocol [20] that we previously adapted for microfluidic cell culture [35]. In brief, to culture hNESC in an OrganoPlate, hNESCs were harvested from wells of a 6 well plate, the harvested hNESC were then re-suspended on Matrigel (catalogue number 354277, lot number 3318549, Discovery Labware, Inc., Two Oak Park, Bedford, MA, USA). 0.7 µL of this Matrigel-cell mix was loaded in assigned chips of the OrganoPlate at a density of 0.03 million cells/µL. After seeding the cells, the plate was loaded into position B3 in the hotel of the Pelican. Afterwards, the plate was moved by the robotic arm to the storage incubator, to start incubation at 37 °C and 5% CO2.

#### Dopaminergic neuronal differentiation

The culture medium preparation “N2B27 medium” consisted of mixed equal amounts of Neurobasal medium (invitrogen/life technologies) and DMEM/F12 medium (invitrogen/life technologies) supplemented with 1% penicillin/streptomycin (life technologies), 2 mM L-glutamine (life technologies), 0.5 X B27 supplement without Vitamin A (life technologies) and 0.5 X N2 supplement (life technologies). The medium to maintain the hNESC in culture “maintenance medium” consisted on N2B27 medium with 0.5 µM PMA (Enzo life sciences), 3 µM CHIR (Axon Medchem) and 150 µM Ascorbic Acid (Sigma Aldrich). The differentiation medium formulation to induce the differentiation of hNESCs towards midbrain dopaminergic neurons “differentiation medium with PMA” consisted in N2B27 medium with 200 µM ascorbic acid, 0.01 ng/µL BDNF (Peprotech), 0.01 ng/µL GDNF (Peprotech), 0.001 ng/µL TGFβ3 (Peprotech), 2.5 µM dbcAMP (Sigma Aldrich) and 1 µM PMA. The function of PMA in this medium preparation was to stimulate the sonic hedghog (SHH) pathway in the cultured hNESCs. Differentiation medium with PMA was changed, every 2 days during the first 6 days of culture in the differentiation process. For the maturation of differentiated neurons PMA was no longer added to the differentiation medium “differentiation medium without PMA” from day 7 onwards, this differentiation medium without PMA was changed every 2 days during 3 weeks. To monitor cellular morphology during differentiation, bright field images were acquired automatically in the Pelican using the microscopy station.

#### Calcium imaging assay

A calcium imaging assay was done on 15 representative chips of the OrganoPlate at day 24 of differentiation. At room temperature, 50 µL of 5 µM cell permeant Fluo-4 AM (Life technologies) in neurobasal medium (Invitrogen/Life technologies) was manually added to the medium inlet well and 20 µL to the medium outlet well of selected chips of the OrganoPlate. Then, the plate was incubated for 30 min at 37°C and 5% CO2. The plate was then placed in an onstage incubator within the microscope. Calcium time-series of spontaneously firing hNESC-derived neurons were then automatically acquired. Images were sampled at a rate of 1 Hz for approximately 5 min, stored as image stacks and analysed using custom Matlab (version 2016b; MathWorks Inc.) scripts. Regions of interest corresponding to individual cells were automatically segmented with an established technique [36] and fluorescence traces were generated for each segmented cell and presented as relative changes in fluorescence intensity ∆*F/F*.

#### Immunofluorescence staining assay

Immunostaining for the dopaminergic neuronal markers class 3 beta tubulin (TUBβIII) and tyrosine hydroxylase (TH), the penultimate enzyme in the biosynthesis of dopamine [20, 50, 51], was performed on representative chips at day 24 of differentiation. Differentiated cells were fixed with 4% paraformaldehyde (PFA) in 1 × phosphate-buffered saline (PBS) for 15 min, by manually adding 70 µL in medium well inlet and 30 µL in medium well outlet followed by permeabilisation with 0.05% Triton-X 100 in 1 × PBS (3 min on ice), and blocking with 10% fetal calf serum (FCS) in 1 × PBS (1h). After washing with 1 × PBS, the primary antibodies mouse anti-TUBβIII (1:2000, Covance) and rabbit anti-TH (1:2000, Santa cruz biotechnology), were incubated for 90 min at room temperature. After washing with 1 × PBS, the secondary antibodies Alexa Fluor 488 Goat Anti-Mouse and Alexa Fluor 568 Goat Anti-Rabbit together with a stain DNA (Hoechst 33342, Invitrogen), were incubated for 2 hours at room temperature. After washing with 1 × PBS and water, confocal images of representative culture chambers were acquired using a confocal microscope (Zeiss LSM 710).

### Automating the differentiation of human neuroepithelial stem cell into dopaminergic neurons

LACS Config Editor was used to develop automated pipelines for the differentiation of hNESC into dopaminergic neurons and time lapse imaging microscopy. The automated pipelines were drafted according to the SiLA communication protocol and command format as previously described [9, 42, 52]. In brief, SiLA uses a Simple Object Access Protocol (SOAP) and a Web Service Description Language (WSDL) documentation, both of which are based on XML. A full library of commands for each device is downloaded once and stored in LACS as a configuration document of the Pelican. LACS Config Editor was used to incorporate the automated pipelines in this configuration file used herein to automate the differentiation of hNESCs into dopaminergic neurons. On the workbench, gel-embedded hNESCs were manually loaded into the culture lanes of a 2-lane OrganoPlate as described above. Then, the plate was put inside the Pelican through Hotel B, and placed inside the storage incubator by the robotic arm. The dispenser was fitted with a 5µL cassette from Biotek and used as a dispensing medium for the media change.

### Statistical Analysis

Three representative chips (n=3) were selected to illustrate the results of this study, in which the statistical analyses were performed by determination of the mean value and the standard deviation of the proportion of dopaminergic neurons within the overall neuronal population.

## Acknowledgements

KIWK, ELM, TH and PV received funding from the SysMedPD project from the European Union’s Horizon 2020 research and innovation program under grant agreement No. 668738. ELM, SH and JJ were also supported by an Aides à la Formation-Recherche training allowance from Fonds National de la Recherche Luxembourg ref. 10099424. The authors thank Miguel Oliveira for his assistance with generation of photographs. The authors also thank Christophe Bouillon for his help to setup the platform.

## Author Contributions Statement

KIWK, RF and ELM proposed and designed the experiments, and wrote the manuscript. JJ and JCS provided the cells. KIWK and ELM run the experiments. KIWK, SH and ELM generated the calcium data and SH analysed the data. MT and MW designed the automated platform. MT, MW and KIWK built the automated platform. All the authors discussed the results and commented on the manuscript.

## Supplementary Information

**Figure S1:**
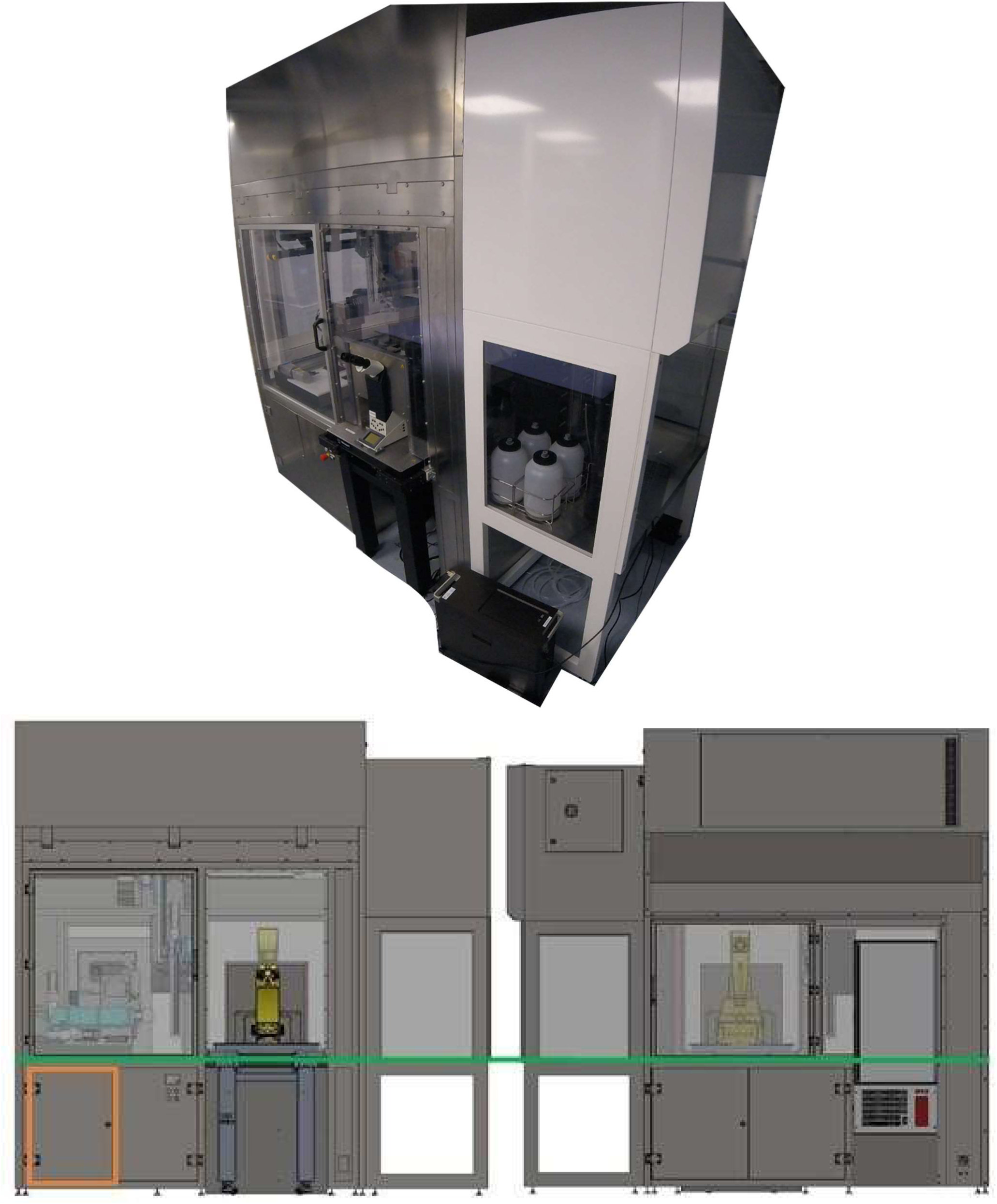
Outside view of automated culture system (top). Front (bottom left) and rear (bottom right) views of Pelican with housing. Yellow (imaging station), light blue (liquid handling station), green (level of stainless steel work surface) and orange (waste containers).

**Figure S2:**
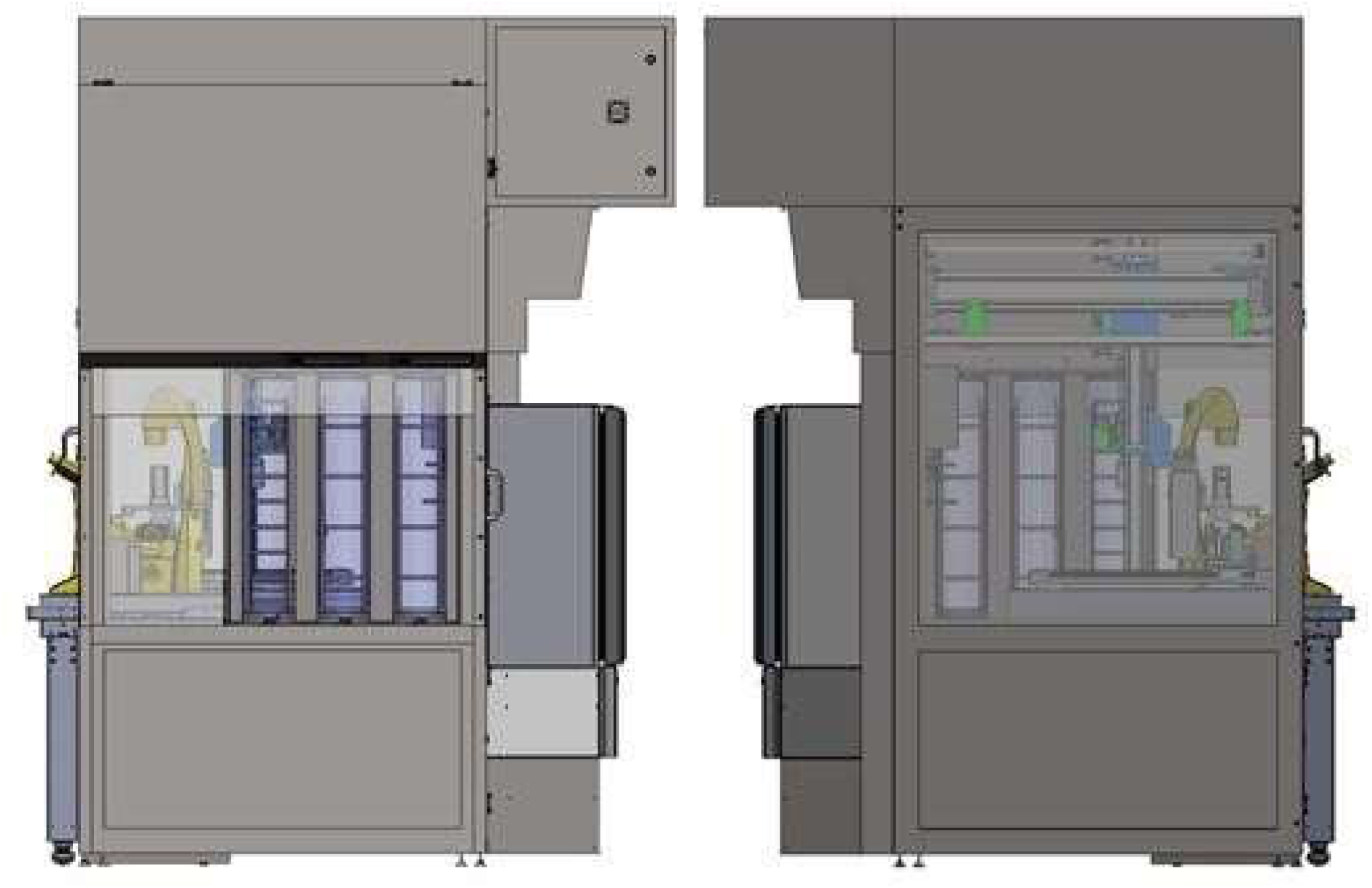
Left and right views of Pelican with housing illustrating the manual work bench and hotels (Left) that connect the Pelican to the outside world and the automated liquid handling station (Right).

**Figure S3:**
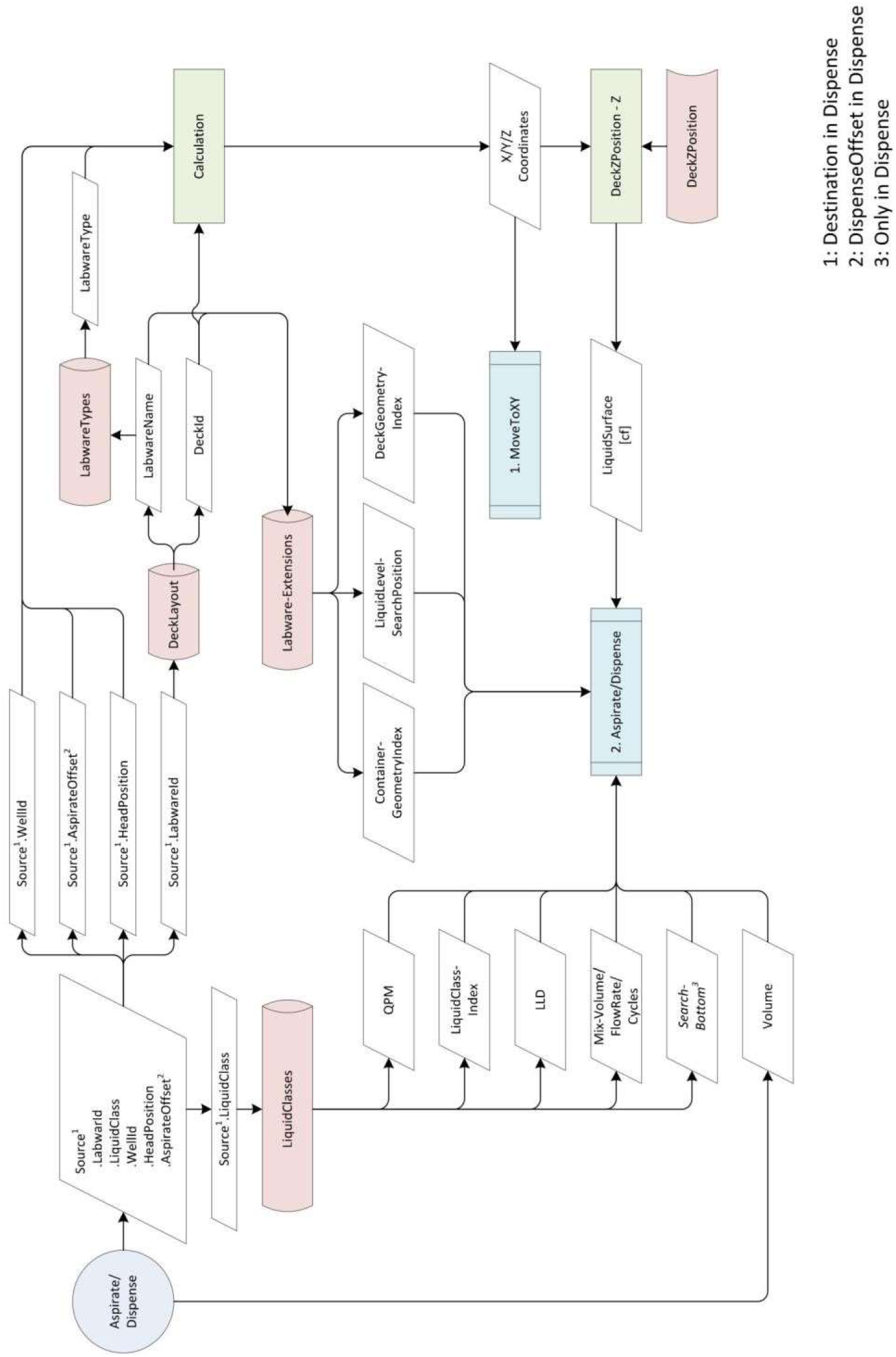
Liquid handler data model for pipetting protocols. Blue (physical action of pipette), red (passive action of pipette: reading from liquid-handler database) and green (passive action of pipette: determining well geometry). QPM=Qualitative Pipette Monitoring. LLD= Liquid Level Detection. The automated pipetting follows this order: pickup a disposable tip, detect the liquid level, aspirate liquid, dispense the liquid into the destination well and finally discard the tip.

## Supplementary Experimental Procedures

### Hardware

#### Housing

The sterile housing of the Pelican (Figure 6B) consists of a stainless steel frame support. The active components of the system, such as the dispenser, pipettor and de-lidder can be flexibly arranged on a perforated stainless steel work surface in any configuration compatible with accessibility via the four-axis gantry robot (Figure 6). This work surface is laterally bordered by an enclosure that guards against manual collision with any robotic moving parts. Large Plexiglas 360º viewing windows enable observation of enclosed processes. The passive components of the Pelican such as the control system, switch cabinets and a variable collection of waste containers (Figure 6B), are housed beneath the work surface.

#### Robot and grippers

A customised four-axis gantry robot (Festo GmbH & Co. KG, Karlsruhe, Germany) was incorporated and mounted at the top of the stainless steel frame support inside the housing. The robot is used to transport any Society for Biomolecular Screening (SBS) format plate (ANSI SLAS 4-2004) and consumables by way of a gripper. The plates were processed at the individual stations, then transported by the robotic arm. The robot and the gripper perform all handling and transport functions of the platform that are not already covered by an individual device. Customised gripper jaws (Fraunhofer IPA, Fraunhofer Society, Stuttgart, Germany) were implemented in order to cover the multitude of plate formats with one gripper. Plates and carriers for consumables can be gripped and stacked by this robot.

#### Liquid-handler

A pipettor transfers various reagents in and out of almost any plate types. Every pipetting step was executed via an air displacement pipetting arm (LiHa-Arm 4x, Hamilton Bonaduz AG, Bonaduz, Switzerland) with disposable tips similar to hand pipettes. Four pipetting units (Z-Excursion Universal Sampler, Hamilton Bonaduz AG, Bonaduz, Switzerland) called ZEUS modules were mounted in a customised X-Y portal (Festo GmbH & Co. KG, Karlsruhe, Germany). Even though the liquid handling robot can move in the X-Y directions, it cannot transport plates. The transport of plates is realised via the four-axis gantry robot. The liquidhandler is equipped with both a capacitive and a pressure liquid level detection system. The capacitive technology is used to detect the liquid level of conductive liquids, while the pressure based technology is for both conductive and non-conductive liquids. Each ZEUS module has an anti-droplet control technology preventing cross contamination from a leaking tip. The pipettor is used to execute precise (pipetting resolution of ±0.1µL) liquid handling steps. It is able to handle volumes from 1µL to 1000µL depending on the available tip sizes in the plant. Depending on volume and the liquid used, contact and non-contact dispensing is possible. The liquid handlers is equipped with 5 positions to hold racks of 96 tips each, a tilting module and 4 heating and cooling stations, the latter 2 are described below. A graphical user interface that follows a specific structure (see Figure S3) was implemented to write all pipetting protocols.

#### Tilter

A Multiflex Tilt Module (Kust, Hamilton Bonaduz AG, Bonaduz, Switzerland) was installed on the liquid handling deck, and is used to tilt SBS plates between a horizontal and a 10° degree incline to completely remove the liquid out of big wells.

#### Heating and cooling positions

Four CPAC Ultraflat & CPAC Microplate Inheco thermoblocks (INHECO Industrial Heating & Cooling GmbH, Planegg, Germany) units, each fitted with a Flat Bottom Adapter (7900016, INHECO Industrial Heating & Cooling GmbH) with positioning frame were installed on the deck of the pipettor as heating and cooling stations. They are used to control the temperature of four SBS plates (4°C-80°C) during time consuming liquid handling protocols. The thermoblocks are controlled by a Multi TEC Control Unit (8900030, INHECO Industrial Heating & Cooling GmbH).

#### Liquid dispenser

A dispenser (EL406, BioTek Instruments Inc, Bad Friedrichshall, Germany) is used to transfer various reagents in and out of SBS plates. It is less precise than the liquid-handler. The liquid dispenser has a dispensing resolution of 2% to 11% depending on the protocol. However, it is much faster than the liquid-handler as it can handle up to 96 wells per step compared to 4 wells per step. For washing steps 96 wells can be filled and emptied simultaneously or with the dispensing unit 8 wells can be filled parallel.

#### Microscopy

A microscope (DMI6000B, Leica AG, Wetzlar, Germany), fitted with an automated onstage stage incubator in an automated stage (SCAN IM 597382, Marzhauser Wetzlar GmbH & Co. KG, Wetzlar, Germany), and a camera (Neo sCMOS 5.5, Andor Technology Ltd, Belfast, Northern Ireland), is used to record images from culture plates. The microscopy assays can be run for a long period of time without perturbing the optimal cell culture conditions because of the presence of the onstage incubator that regulates the CO_2_ and the temperature levels.

#### Onstage incubator

A small incubator housing (H301-K-Frame, Okolab SRL, Naples, Italy) was installed on the stage of the microscope. This incubator was used to store plates in SBS format under regulated conditions. The temperature and CO_2_ of the onstage incubator is controlled during microscopic examination by a temperature controller (H301-T-UNIT-BL-PLUS Boldline, Okolab SRL, Naples, Italy) and a CO_2_ controller (0506.000, Pecon GmbH, Erbach, Germany) respectively.

#### Storage Incubator

An incubator (STX-44, LiCONiC AG, Mauren, Principality of Liechtenstein) is used to store plates in SBS format under regulated conditions in terms of temperature, CO_2_ and humidity. A varying number of plates, of varying height, may be stored depending on the format of the racks within the incubator. Two standard Liconic stackers, each with a pitch of 23mm suitable for plates with a maximum height of 17mm, were mounted inside the incubator. Each stacker can store up to 22 plates.

#### de-lidder

Two de-lidders are used to remove the lids of up to two SBS plates located in the process at any given time. The lids are kept by the de-lidder while the plates are being processed. Another function of this component is to place the lids back on after the plates were processed. It has an internal pressure-based check that verifies the correct handling of the lids for removal and closure.

#### Hotel

The hotels are used as a secure transfer station between the outside world and inside the housing. All consumables and culture plates are fed in and discharged via the hotels. There are 3 hotels (A, B, and C). Hotel A and hotel B are intended for the loading and discharge of pipette tip racks. Pipette tips may be incorporated on four racks each with 96 tips. Plates in SBS format may be fed in and discharged in all hotels.

#### Waste Handling

A waste position on the liquid handling deck is available for tip disposal while running a liquid handling process. Pipette tips exit via a downpipe under the work surface where they are dropped into a collection container, which can easily be pulled forward on a rail between the switch cabinets (Figure 6B). Media waste is collected in SBS plates, and removed through the hotels.

### Environmental control

#### Sterility

All the components of the Pelican were contained within an enclosure housing as described above. Air flows at a rate of pressure 161 Pa and wind speed of 0.45 m/s through a filter at the top of the automation enclosure, with a continuous laminar flow down through the perforated work surface, except where devices deviate the air flow. This air flow is designed to maintain sterility (biosafety level 1) of the automation enclosure.

#### Temperature

The temperature levels were controlled on most positions of the Pelican. The temperature was regulated through the four heating and cooling positions (4°C-80°C) while changing the media on the liquidhandler deck. It was also controlled on the stage incubator (25°C-45°C), while acquiring images, and the incubator is always kept at 37°C at all times.

#### Carbon dioxide

The Pelican was fed with pure carbon dioxide (99.99%) at a pressure of 1.5bar. The carbon dioxide levels inside the onstage incubator was regulated (0-7%) and the storage incubator was kept at 5% CO_2_at all times.

#### Humidity

While plates were stored inside the storage incubator, the relative humidity was kept at 95%.

#### Practicability

A user manual details the procedure for operation of the Pelican Cell culture Observatory (available upon request). This document detailed not only how to operate the system, but also how to extend the current setup to include more devices, and how to integrate the said devices in terms of hardware and software development.

## References

[1] Dunn DA, Feygin I. Challenges and solutions to ultra-high-throughput screening assay miniaturization: submicroliter fluid handling. Drug Discovery Today. 2000 Dec;5(12, Supplement 1):84–91. Available from: http://www.sciencedirect.com/science/article/pii/S1359644600000647.

[2] Kong F, Yuan L, Zheng YF, Chen W. Automatic Liquid Handling for Life Science: A Critical Review of the Current State of the Art. J Lab Autom. 2012 Jun;17(3):169–185. Available from: http://dx.doi.org/10.1177/2211068211435302.

[3] Dauwalder O, Landrieve L, Laurent F, de Montclos M, Vandenesch F, Lina G. Does bacteriology laboratory automation reduce time to results and increase quality management? Clinical Microbiology and Infection. 2016 Mar;22(3):236–243. Available from: http://www.sciencedirect.com/science/article/pii/S1198743X15009787.

[4] Arrasate M, Finkbeiner S. Automated microscope system for determining factors that predict neuronal fate. Proc Natl Acad Sci U S A. 2005 Mar;102(10):3840–3845. Available from: http://www.ncbi.nlm.nih.gov/pmc/articles/PMC553329/.

[5] Skibinski G, Finkbeiner S. Longitudinal measures of proteostasis in live neurons: Features that determine fate in models of neurodegenerative disease. FEBS Letters. 2013 Apr;587(8):1139–1146. Available from: http://onlinelibrary.wiley.com/doi/10.1016/j.febslet.2013.02.043/abstract.

[6] Triaud F, Clenet DH, Cariou Y, Le Neel T, Morin D, Truchaud A. Evaluation of Automated Cell Culture Incubators. JALA: Journal of the Association for Laboratory Automation. 2003 Dec;8(6):82–86. Available from: http://journals.sagepub.com/doi/abs/10.1016/s1535-5535%2803%2900018-2.

[7] Halldorsson S, Lucumi E, Gómez-Sjöberg R, Fleming RMT. Advantages and challenges of microfluidic cell culture in polydimethylsiloxane devices. Biosensors and Bioelectronics. 2015 Jan;63:218–231. 00126. Available from: http://www.sciencedirect.com/science/article/pii/S0956566314005302.

[8] Carvalho MC. Integration of Analytical Instruments with Computer Scripting. Journal of Laboratory Automation. 2013 Aug;18(4):328–333. 00001 PMID: 23413273. Available from: http://jla.sagepub.com/content/18/4/328.

[9] Bär H, Hochstrasser R, Papenfub B. SiLA: Basic standards for rapid integration in laboratory automation. J Lab Autom. 2012 Apr;17(2):86–95.

[10] Lees AJ, Hardy J, Revesz T. Parkinson’s disease. The Lancet. 2009 Jun;373(9680):2055–2066. Available from: http://www.sciencedirect.com/science/article/pii/S014067360960492X.

[11] Abdullah R, Basak I, Patil KS, Alves G, Larsen JP, Møller SG. Parkinson’s disease and age: The obvious but largely unexplored link. Experimental Gerontology. 2015 Aug;68:33–38. Available from: http://www.sciencedirect.com/science/article/pii/S053155651400271X.

[12] Surmeier DJ, Schumacker PT. Calcium, Bioenergetics, and Neuronal Vulnerability in Parkinson’s Disease. J Biol Chem. 2013 Apr;288(15):10736–10741. Available from: http://www.jbc.org/content/288/15/10736.

[13] Bellucci A, Mercuri NB, Venneri A, Faustini G, Longhena F, Pizzi M, et al. Review: Parkinson’s disease: from synaptic loss to connectome dysfunction. Neuropathol Appl Neurobiol. 2016 Feb;42(1):77–94. Available from: http://onlinelibrary.wiley.com/doi/10.1111/nan.12297/abstract.

[14] Chinta SJ, Andersen JK. Dopaminergic neurons. The International Journal of Biochemistry & Cell Biology. 2005 May;37(5):942–946. 00102. Available from: http://www.sciencedirect.com/science/article/pii/S1357272504003711.

[15] Schöndorf DC, Aureli M, McAllister FE, Hindley CJ, Mayer F, Schmid B, et al. iPSC-derived neurons from GBA1-associated Parkinson’s disease patients show autophagic defects and impaired calcium homeostasis. Nat Commun. 2014 Jun;5. 00002. Available from: http://www.nature.com/ncomms/2014/140606/ncomms5028/full/ncomms5028.html.

[16] Muñoz P, Huenchuguala S, Paris I, Segura-Aguilar J. Dopamine Oxidation and Autophagy. Parkinsons Dis. 2012;2012. 00000. Available from: http://www.ncbi.nlm.nih.gov/pmc/articles/PMC3433151/.

[17] Pfeiffer RF, Wszolek ZK, Ebadi M. Parkinson’s Disease, Second Edition. CRC Press; 2012. Available from: https://www.crcpress.com/Parkinsons-Disease-Second-Edition/Pfeiffer-Wszolek-Ebadi/p/book/9781439807149.

[18] Hillje AL, Schwamborn JC. Utilization of stem cells to model Parkinson’s disease – current state and future challenges. Future Neurology. 2016 Apr;11(2):171–186. Available from: http://www.futuremedicine.com/doi/abs/10.2217/fnl.16.7.

[19] Takahashi K, Tanabe K, Ohnuki M, Narita M, Ichisaka T, Tomoda K, et al. Induction of Pluripotent Stem Cells from Adult Human Fibroblasts by Defined Factors. Cell. 2007 Nov;131(5):861–872. 08307. Available from: http://www.sciencedirect.com/science/article/pii/S0092867407014717.

[20] Reinhardt P, Glatza M, Hemmer K, Tsytsyura Y, Thiel CS, Höing S, et al. Derivation and Expansion Using Only Small Molecules of Human Neural Progenitors for Neurodegenerative Disease Modeling. PLoS ONE. 2013 Mar;8(3):e59252. 00023. Available from: http://dx.plos.org/10.1371/journal.pone.0059252.

[21] Gómez-Sjöberg R, Leyrat AA, Pirone DM, Chen CS, Quake SR. Versatile, Fully Automated, Microfluidic Cell Culture System. Analytical Chemistry. 2007 Nov;79(22):8557–8563. 00446. Available from: http://pubs.acs.org/doi/abs/10.1021/ac071311w.

[22] Lecault V, Vaninsberghe M, Sekulovic S, Knapp D, Wohrer S, Bowden W, et al. High-throughput analysis of single hematopoietic stem cell proliferation in microfluidic cell culture arrays. Nat Methods. 2011 Jul;8(7):581–586. 00174. Available from: http://www.hubmed.org/display.cgi?uids=21602799.

[23] Croushore C, Supharoek S, Lee C, Jakmunee J, Sweedler J. Microfluidic device for the selective chemical stimulation of neurons and characterization of peptide release with mass spectrometry. Anal Chem. 2012 Nov;84(21):9446–9452. Available from: http://www.hubmed.org/display.cgi?uids=23004687.

[24] Shintu L, Baudoin R, Navratil V, Prot J, Pontoizeau C, Defernez M, et al. Metabolomics-on-a-chip and predictive systems toxicology in microfluidic bioartificial organs. Anal Chem. 2012 Feb;84(4):1840–1848. 00053. Available from: http://www.hubmed.org/display.cgi?uids=22242722.

[25] Oedit A, Vulto P, Ramautar R, Lindenburg PW, Hankemeier T. Lab-on-a-Chip hyphenation with mass spectrometry: strategies for bioanalytical applications. Current Opinion in Biotechnology. 2015 Feb;31:79–85. 00000. Available from: http://www.sciencedirect.com/science/article/pii/S0958166914001517.

[26] Huh D, Matthews BD, Mammoto A, Montoya-Zavala M, Hsin HY, Ingber DE. Reconstituting Organ-Level Lung Functions on a Chip. Science. 2010 Jun;328(5986):1662–1668. Available from: http://science.sciencemag.org/content/328/5986/1662.

[27] Vulto P, Podszun S, Meyer P, Hermann C, Manz A, Urban GA. Phaseguides: a paradigm shift in microfluidic priming and emptying. Lab Chip. 2011 May;11(9):1596–1602. 00031. Available from: http://pubs.rsc.org/en/content/articlelanding/2011/lc/c0lc00643b.

[28] Trietsch SJ, Israëls GD, Joore J, Hankemeier T, Vulto P. Microfluidic titer plate for stratified 3D cell culture. Lab on a Chip. 2013;13(18):3548. Available from: http://xlink.rsc.org/?DOI=c3lc50210d.

[29] van Duinen V, Trietsch SJ, Joore J, Vulto P, Hankemeier T. Microfluidic 3D cell culture: from tools to tissue models. Current Opinion in Biotechnology. 2015 Dec;35:118–126. Available from: http://www.sciencedirect.com/science/article/pii/S0958166915000713.

[30] Junaid A, Mashaghi A, Hankemeier T, Vulto P. An end-user perspective on Organ-on-a-Chip: Assays and usability aspects. Current Opinion in Biomedical Engineering. 2017 Mar;1:15–22. Available from: http://www.sciencedirect.com/science/article/pii/S246845111730003X.

[31] Grossmann G, Guo W, Ehrhardt D, Frommer W, Sit R, Quake S, et al. The RootChip: an integrated microfluidic chip for plant science. Plant Cell. 2011 Dec;23(12):4234–4240. Available from: http://www.hubmed.org/display.cgi?uids=22186371.

[32] Antia M, Herricks T, Rathod P. Microfluidic modeling of cell-cell interactions in malaria pathogenesis. PLoS Pathog. 2007 Jul;3(7):0939–0948. 00064. Available from: http://www.hubmed.org/display.cgi?uids=17658948.

[33] Ho SY, Chao CY, Huang HL, Chiu TW, Charoenkwan P, Hwang E. NeurphologyJ: An automatic neuronal morphology quantification method and its application in pharmacological discovery. BMC Bioinformatics. 2011 Jun;12:230. Available from: http://www.ncbi.nlm.nih.gov/pmc/articles/PMC3121649/.

[34] Vedel S, Tay S, Johnston DM, Bruus H, Quake SR. Migration of cells in a social context. Proceedings of the National Academy of Sciences. 2013 Jan;110(1):129–134. Available from: http://www.pnas.org/cgi/doi/10.1073/pnas.1204291110.

[35] Lucumi Moreno E, Hachi S, Hemmer K, Trietsch SJ, Baumuratov AS, Hankemeier T, et al. Differentiation of neuroepithelial stem cells into functional dopaminergic neurons in 3D microfluidic cell culture. Lab Chip. 2015 May;15(11):2419–2428. Available from: http://pubs.rsc.org/en/content/articlelanding/2015/lc/c5lc00180c.

[36] Diego F, Reichinnek S, Both M, Hamprecht FA. Automated identification of neuronal activity from calcium imaging by sparse dictionary learning. In: 2013 IEEE 10th International Symposium on Biomedical Imaging (ISBI); 2013. p. 1058–1061.

[37] Piracci A. Advantages of Non-Contact Dispensing in SMT Assembly Processes. In: SMTA International Conference Proceedings; 2000. Available from: http://www.smta.org/knowledge/proceedings_abstract.cfm?PROC_ID=619.

[38] American Society for Testing and Material. Standard Specification for Transferring Information Between Clinical Instruments and Computer Systems (Withdrawn 2002). In: In Annual Book of ASTM Standards. vol. 14.01. West Conshohocken, PA; 1997. Available from: https://www.astm.org/DATABASE.CART/WITHDRAWN/E1394.htm.

[39] Smith B, Ceusters W. HL7 RIM: an incoherent standard. Stud Health Technol Inform. 2006;124:133–138.

[40] Joshi S, Pillai R. LECIS Commentary; 2002. Available from: https://www.ergotech.com/lecis.org/documents/UserSpace/LECIS_commentary.pdf.

[41] Roth A, Jopp R, Schäfer R, Kramer GW. Automated Generation of Animl Documents by Analytical Instruments. JALA: Journal of the Association for Laboratory Automation. 2006 Aug;11(4):247–253. Available from: http://journals.sagepub.com/doi/abs/10.1016/j.jala.2006.05.013.

[42] Bäar H, Syré U. JALA: Journal of the Association for Infoteam SiLA Library Simplifies Device Integration. Laboratory Automation. 2011 Oct;16(5):371–376. Available from: http://journals.sagepub.com/doi/abs/10.1016/j.jala.2011.05.003.

[43] Paull D, Sevilla A, Zhou H, Hahn AK, Kim H, Napolitano C, et al. Automated, high-throughput derivation, characterization and differentiation of induced pluripotent stem cells. Nat Meth. 2015 Sep;12(9):885–892. Available from: https://www.nature.com/nmeth/journal/v12/n9/full/nmeth.3507.html.

[44] Konagaya S, Ando T, Yamauchi T, Suemori H, Iwata H. Long-term maintenance of human induced pluripotent stem cells by automated cell culture system. Scientific Reports. 2015 Nov;5:srep16647. Available from: https://www.nature.com/articles/srep16647.

[45] Soares FAC, Chandra A, Thomas RJ, Pedersen RA, Vallier L, Williams DJ. Investigating the feasibility of scale up and automation of human induced pluripotent stem cells cultured in aggregates in feeder free conditions. Journal of Biotechnology. 2014 Mar;173(Supplement C):53–58. Available from: http://www.sciencedirect.com/science/article/pii/S016816561300552X.

[46] Kami D, Watakabe K, Yamazaki-Inoue M, Minami K, Kitani T, Itakura Y, et al. Large-scale cell production of stem cells for clinical application using the automated cell processing machine. BMC Biotechnology. 2013 Nov;13:102. Available from: https://doi.org/10.1186/1472-6750-13-102.

[47] Thomas RJ, Anderson D, Chandra A, Smith NM, Young LE, Williams D, et al. Automated, scalable culture of human embryonic stem cells in feeder-free conditions. Biotechnol Bioeng. 2009 Apr;102(6):1636–1644. Available from: http://onlinelibrary.wiley.com/doi/10.1002/bit.22187/abstract.

[48] Terstegge S, Laufenberg I, Pochert J, Schenk S, Itskovitz-Eldor J, Endl E, et al. Automated maintenance of embryonic stem cell cultures. Biotechnol Bioeng. 2007 Jan;96(1):195–201. Available from: http://onlinelibrary.wiley.com/doi/10.1002/bit.21061/abstract.

[49] Exner N, Lutz AK, Haass C, Winklhofer KF. Mitochondrial dysfunction in Parkinson’s disease: molecular mechanisms and pathophysiological consequences. EMBO J. 2012 Jul;31(14):3038–3062. 00114 PMID: 22735187.

[50] Daadi MM, Grueter BA, Malenka RC, Jr DER, Steinberg GK. Dopaminergic Neurons from Midbrain-Specified Human Embryonic Stem Cell-Derived Neural Stem Cells Engrafted in a Monkey Model of Parkinson’s Disease. PLOS ONE. 2012 Jul;7(7):e41120. Available from: http://journals.plos.org/plosone/article?id=10.1371/journal.pone.0041120.

[51] Yan Y, Yang D, Zarnowska ED, Du Z, Werbel B, Valliere C, et al. Directed Differentiation of Dopaminergic Neuronal Subtypes from Human Embryonic Stem Cells. Stem Cells. 2005 Jun;23(6):781–790. 00390. Available from: http://doi.wiley.com/10.1634/stemcells.2004-0365.

[52] McIntosh RL, Yau A. A Flexible and Robust Peer-to-Peer Architecture with XML-Based Open Communication for Laboratory Automation. JALA: Journal of the Association for Laboratory Automation. 2003 Feb;8(1):38–45. Available from: http://journals.sagepub.com/doi/abs/10.1016/S1535-5535-04-00240-0.

